# Screening in serum-derived medium reveals differential response to compounds targeting metabolism

**DOI:** 10.1101/2023.02.25.529972

**Authors:** Keene L. Abbott, Ahmed Ali, Dominick Casalena, Brian T. Do, Raphael Ferreira, Jaime H. Cheah, Christian K. Soule, Amy Deik, Tenzin Kunchok, Daniel R. Schmidt, Steffen Renner, Sophie E. Honeder, Michelle Wu, Sze Ham Chan, Tenzin Tseyang, Daniel Greaves, Peggy P. Hsu, Christopher W. Ng, Chelsea J. Zhang, Ali Farsidjani, Iva Monique T. Gramatikov, Nicholas J. Matheson, Caroline A. Lewis, Clary B. Clish, Matthew G. Rees, Jennifer A. Roth, Lesley Mathews Griner, Alexander Muir, Douglas S. Auld, Matthew G. Vander Heiden

## Abstract

A challenge for screening new candidate drugs to treat cancer is that efficacy in cell culture models is not always predictive of efficacy in patients. One limitation of standard cell culture is a reliance on non-physiological nutrient levels to propagate cells. Which nutrients are available can influence how cancer cells use metabolism to proliferate and impact sensitivity to some drugs, but a general assessment of how physiological nutrients affect cancer cell response to small molecule therapies is lacking. To enable screening of compounds to determine how the nutrient environment impacts drug efficacy, we developed a serum-derived culture medium that supports the proliferation of diverse cancer cell lines and is amenable to high-throughput screening. We used this system to screen several small molecule libraries and found that compounds targeting metabolic enzymes were enriched as having differential efficacy in standard compared to serum-derived medium. We exploited the differences in nutrient levels between each medium to understand why medium conditions affected the response of cells to some compounds, illustrating how this approach can be used to screen potential therapeutics and understand how their efficacy is modified by available nutrients.

## INTRODUCTION

Studies of cancer cells in standard culture conditions have long been used as a tractable tool for drug discovery. Cell culture provides unparalleled experimental flexibility, scalability, and low cost to identify and understand the response to cancer therapeutics; however, drug responses in culture are not always predictive of drug response in animal models or in patients^1–6^. A limitation of standard culture systems that may contribute to this phenomenon is a failure to model many of the cell-extrinsic factors that influence tumor growth and progression, including cancer cell interactions with immune cells^7,8^, stromal cells^9–11^, and the extracellular matrix^12–14^. Additionally, it is becoming increasingly apparent that nutrient availability can influence the response to some drugs^1–5,15,16^. For example, most cancer cells are sensitive to inhibitors of the metabolic enzyme glutaminase when cultured in standard media^2,4,6^, but become resistant to glutaminase inhibitors when cultured in serum-like media^4^ or when grown as tumors in mice^6^. Similarly, cancer cells were found to be protected from 5-fluorouracil efficacy when cultured in the presence of physiologically relevant levels of the purine catabolism end product uric acid^5^.

Culturing cancer cells in media containing more physiological levels of defined nutrients is an approach that is tractable and has been shown to affect metabolism and alter sensitivity to some drugs in a manner that may better reflect the responses of tumors in mice^4,5,15,17,18^. However, while synthetic culture media have the advantage of nutrients being supplied at known concentrations, they lack the biological complexity of nutrients present in blood and tissues. This includes the absence of many lipid species and undefined low abundance metabolites. An alternative approach for culturing cells in more physiological nutrient levels is the use of biological fluids as a culture medium^4,19–21^, since these fluids contain all nutrients present in the animal. Additionally, culturing cells in animal-derived serum is cost effective, and has recently been used to understand mechanisms of drug sensitivity and resistance to glutaminase inhibitors^4^. The biological complexity of nutrients present in serum may be of particular interest for phenotypic drug screening to identify compounds with anticancer activity, as some may target pathways involving micronutrients or lipid metabolism that are less well modeled in defined media.

Screening approaches to find candidate compounds whose efficacy is impacted by available nutrients could complement existing screens using standard media, and might identify hits that would otherwise be masked by non-physiological nutrient levels. In this study, we optimize a system for screening in serum-based conditions that is amenable for culture of >482 cancer cell lines and allows for testing differential drug sensitivity in a high-throughput format. Using this system to screen several small molecule libraries, we find that inhibitors of metabolic proteins show differential efficacy between standard and more physiologic nutrient conditions, and illustrate how these differences may be explained mechanistically. We therefore demonstrate how the use of a serum-based medium may be utilized as a tool for drug discovery.

## RESULTS

### Characterization of a serum-derived culture medium optimized for high-throughput screening

To facilitate high-throughput compound screening in cells cultured in serum-based media conditions, we first optimized ABS to allow for direct cell plating and subsequent growth in 1536-well plates. As previously observed^4^, cancer cells seeded directly into culture plates in 100% ABS exhibited poor attachment; additionally, we found dispensing ABS through automated reagent dispensers led to clogging of those systems. We hypothesized that high protein content in ABS contributed to low cell attachment^22^. Therefore, we performed ultrafiltration with a 10 kDa filter to reduce protein content and collected the small molecule-containing filtrate, which we termed basal flow-through ABS (ftABS^B^) **(Figure S1A, B).** To assess how levels of metabolites were altered due to ultrafiltration, we performed quantitative metabolomics^23^ as well as untargeted metabolomics of ABS and ftABS^B^. ftABS^B^ maintained most polar metabolites at similar levels to those found in 100% ABS **(Figure 1A, S1C-D, and Table S1).** Additionally, while some lipid species were lost during ultrafiltration, many were retained including some that are not present in synthetic media **(Figure S1E and Table S1).** After supplementing the ABS ultrafiltrate with 10% dialyzed FBS as a growth factor source, we obtained a media we termed ftABS^B+F^ that allowed cancer cells to attach to culture dishes when directly plated using automated reagent dispensers.

**Figure 1.**
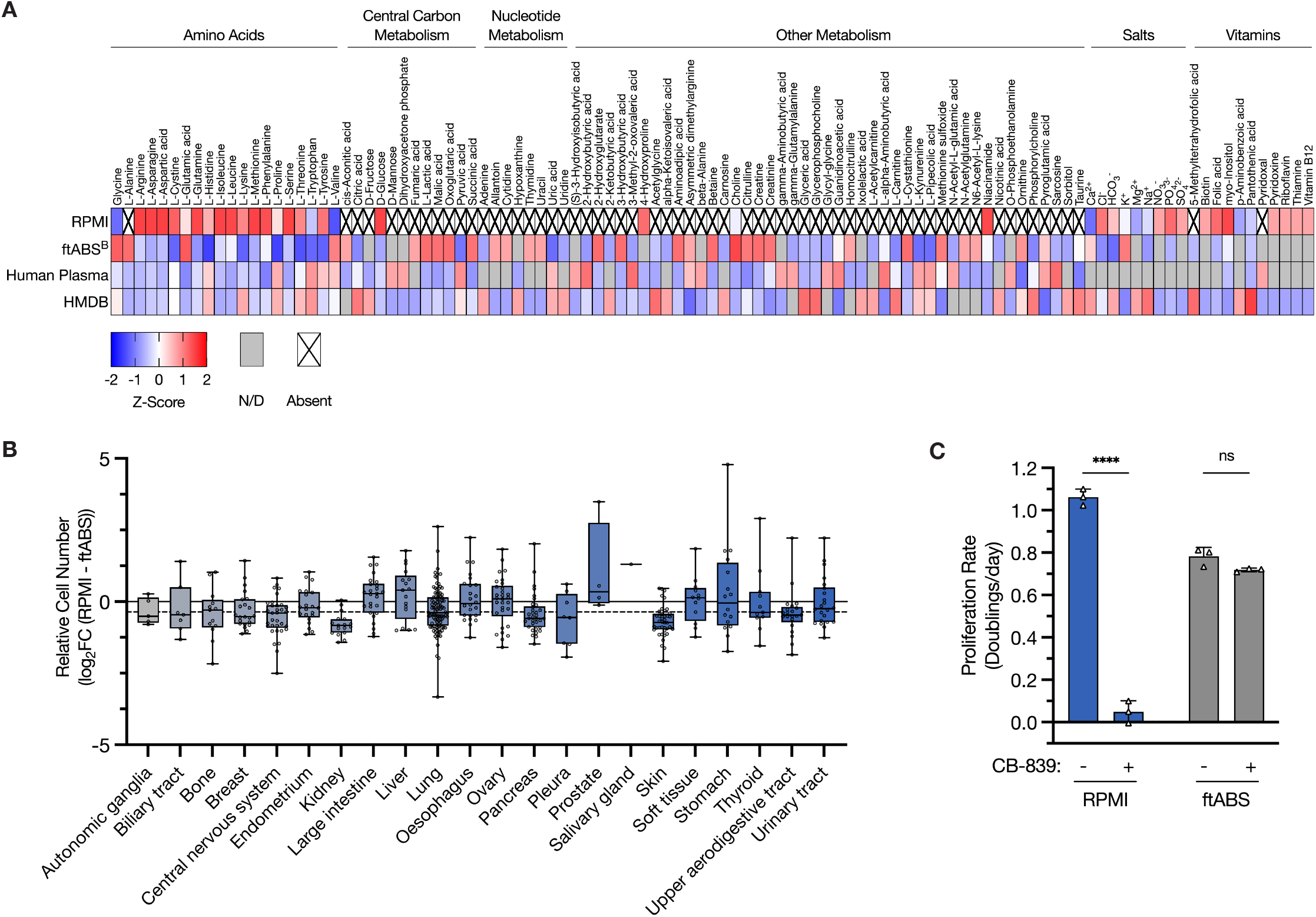
Characterization of a serum-derived culture medium optimized for high-throughput screening. **(A)** Heatmap depicting relative concentrations of the indicated components present in RPMI, measured in basal flow-through adult bovine serum (ftABS^B^), measured in human plasma, or reported in the Human Metabolome Database (HMDB). Data presented within each column were z-score normalized. ND: not determined or not detected in the samples, or values not reported on HMDB. Absent: not present in RPMI. See Table S1 for heatmap values and metabolite concentrations. **(B)** log_2_ fold change in relative cell number for 482 barcoded cancer cell lines grouped by tissue of origin when cultured in ftABS compared to RPMI for 5 days. The median (box center line), interquartile range (IQR) (box) and 1.5×10R (whiskers) are plotted; the dashed line represents the population median. **(C)** Proliferation rate of A549 cells in cultured in RPMI or ftABS and treated with or without CB-839 for 72 hr. Data shown are mean ± SD from three technical replicates. Statistical test was performed using multiple unpaired t test (ns, not significant; ****p < 0.0001).

When culturing A549 cells in ftABS^B+F^, we observed that cells proliferated slower in this medium than they did in RPMI **(Figure S1F).** Consistent with previous measurements^4^, cystine is present at sub-physiologic levels in both ABS and ftABS^B^ (as compared to 25-100 μM in adult human plasma^24,25^, **Figure 1A and Table S1).** Restoring cystine in ftABS^B+F^ to a more physiologically relevant concentration of 25 μM resulted in a medium we termed ftABS^B+FC^ that supported proliferation of A549 cells at rates that approached those observed in RPMI **(Figure S1F).**

Despite cystine addition, cells cultured in ftABS^B+FC^ stopped proliferating after two days unless the medium was refreshed daily **(Figure S1G),** suggesting one or more metabolites may become depleted below a critical threshold over time in culture. We examined whether daily media changes were feasible for high-throughput screening, but observed variable cell losses in 1536-well plates with media replacement, regardless of whether various approaches were used to promote improved cell attachment **(Figure S1H).** Thus, to overcome this limitation and allow screening periods of more than 2 days, we next considered which metabolites may become limiting for cell proliferation in ftABS, or if the ratio of culture volume to cell number could be adjusted to prevent metabolite depletion over a longer assay period. Glutamine is avidly consumed by cancer cells in vitro^11^, and given its sub-physiologic concentration of 100 μM in ftABS^B^ (as compared to ~500 μM in adult human plasma^24,25^, **Figure 1A and Table S1),** we hypothesized that it may be limiting for proliferation. To test this hypothesis, and to identify other nutrients that may become limiting, we analyzed metabolite consumption by A549 cells cultured in different volumes of RPMI or ftABS^B+FC^ and found that glutamine was indeed depleted or nearly depleted after three days **(Figure S2A).** Also consistent with nutrient depletion in ftABS, the falloff in proliferation rate of A549 cells in ftABS^B+FC^ over time could be suppressed by culture in a larger volume of media **(Figure S2B).** We therefore supplemented ftABS^B+FC^ with an additional 400 μM glutamine to more closely match the human physiologic range, and found that it improved stable rates of cell proliferation over a longer period of time without media change **(Figure S1I).** This final, complete version of media we used for screening is henceforth referred to as ftABS, and we further optimized the cell plating density to allow for exponential cell proliferation over the course of three days in ftABS medium **(Figure S1J).**

We next tested whether ftABS could be used as a culture medium to screen diverse cancer cell lines. Utilizing the PRISM screening platform, in which 482 barcoded cancer cell lines are co-cultured over several days^26^, we found that most cell lines display a similar relative proliferation when cultured in ftABS compared to RPMI **(Figure 1B and Table S2).** We also assessed the proliferation rate of individual human and murine cancer cells and found that a diverse set of cell lines all proliferated in ftABS, in many cases exhibiting minimal differences in proliferation rates when cultured in ftABS compared to RPMI **(Figure S2C).** These data indicate that ftABS contains metabolites at sufficient levels to support the proliferation of many cancer cell lines from diverse tissues of origin over a time period amenable to high-throughput screening.

We next verified that culturing cells in ftABS recapitulates the reduced sensitivity to the glutaminase inhibitor CB-839 that was previously observed when cells were cultured in 100% ABS^4^, providing a positive control for differential drug sensitivity in screens involving ftABS compared to standard culture medium. Indeed, A549 cells treated with CB-839 were sensitive to this inhibitor when cultured in RPMI but were insensitive when cultured in ftABS **(Figure 1C).** Altogether, these data demonstrate that ftABS is a serum-based culture medium that supports the growth of most cancer cell lines and allows for high-throughput compound screening.

### Compounds targeting metabolism are differentially enriched in cells cultured in standard versus serum-derived medium

Next, we investigated how culturing A549 cells in RPMI versus ftABS affects response to a drug screening library containing 3,491 compounds with annotated mechanisms of action^27^. We found that the efficacy of most compounds in the library was highly correlated for cells cultured in RPMI compared to ftABS **(Figure 2A),** indicating that the effects of most compounds in this library are not altered by environmental nutrient levels. Of the compounds with differential efficacy in the screen, we noted many of them were annotated to target metabolic enzymes. To assess whether compounds targeting metabolism are enriched as having differential efficacy between the two media conditions, we generated a consensus metabolic library gene list by intersecting three metabolic gene lists either generated in-house or from the literature^28,29^ **(Table S3).** Interestingly, we found that many compounds targeting metabolic proteins were particularly enriched for being more effective at reducing cell proliferation in RPMI **(Figure 2A-B).** We performed a similar analysis by generating a consensus gene list comprising signaling proteins from KEGG **(Table S3),** and found that compounds targeting signaling proteins were not enriched as having differential efficacy across the two media conditions **(Figure 2C).**

**Figure 2.**
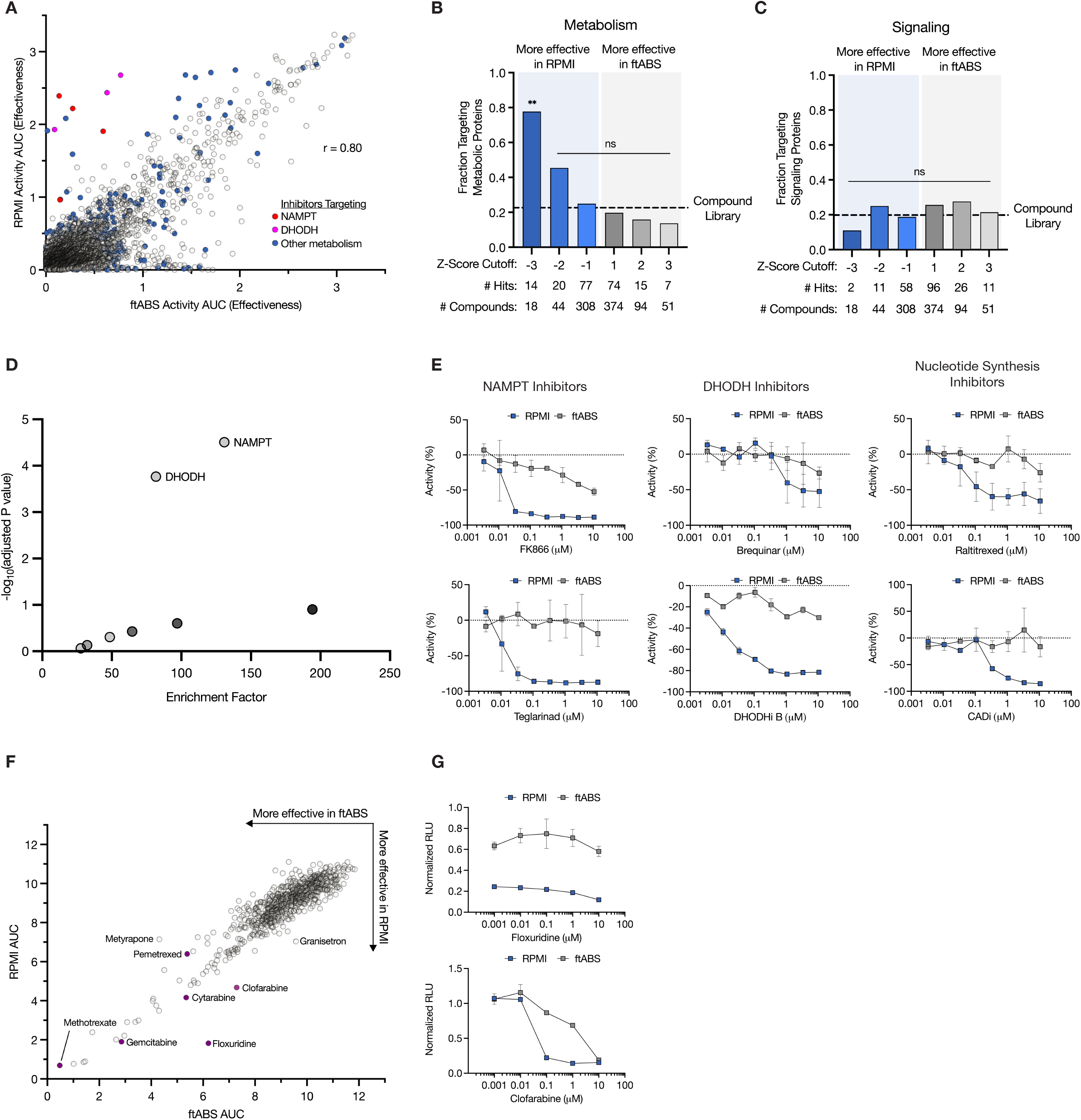
Compounds targeting metabolic enzymes show differential efficacy in standard versus serum-derived medium. **(A)** Plot of activity area under the curve (AUC) for A549 cells cultured in RPMI or ftABS and treated with the Novartis Institute of BioMedical Research Mechanism of Action Box (MoA Box) compound library. Cells were treated for 72 hr with 8 doses of each drug up to a maximum dose of 10.7 μM, and dose response used to calculate AUC. Drugs in red are NAMPT inhibitors, drugs in magenta are DHODH inhibitors, and drugs in blue are inhibitors of other metabolic enzymes. **(B-C)** Analysis of whether compounds targeting metabolic enzymes (B) or signaling proteins (C)were differentially active in the two media used in the screen shown in Figure 2A. Each compound included in the MoA Box compound library was classified based on whether or not it targeted a metabolic enzyme or signaling protein and then binned according to z-score based on the difference in AUC between ftABS and RPMI. Statistical tests were performed using Fisher’s exact test (ns, not significant; **p < 0.01). **(D)** Drug enrichment analysis of compounds from the MoA Box screen shown in Figure 2A. The enrichment factor is a score that represents the fraction of compounds targeting each gene that are differentially active between RPMI or ftABS. **(E)** Dose-response curves of A549 cells treated with the indicated compounds targeting NAMPT, DHODH, or other enzymes involved in nucleotide synthesis from the screen shown in Figure 2A. Data shown are mean ± SD from two technical replicates. **(F)** Plot of AUC for A549 cells cultured in RPMI or ftABS and treated with the SCREEN-WELL^®^ FDA v. 2.0 Approved Drug Library. Cells were treated for 72 hr with 5 doses of each drug up to a maximum dose of 10 μM, and dose response used to calculate AUC. Drugs in purple are nucleotide synthesis inhibitors. **(G)** Dose-response curves of A549 cells treated with the indicated compounds targeting nucleotide synthesis from the screen shown in 2F. Data shown are mean ± SD from two technical replicates.

We next performed enrichment factor analysis of significant screen hits and found that compounds targeting the metabolic enzymes NAMPT, which is involved in NAD^+^ salvage, and DHODH, which is required for pyrimidine synthesis, were particularly effective in impairing cell proliferation in RPMI relative to ftABS **(Figure 2D-E).** Additional compounds targeting nucleotide synthesis were also enriched as being more effective in RPMI than in ftABS **(Figure 2E).** To expand this analysis to more compounds with known mechanisms of action, we also screened a small library containing FDA-approved compounds **(Figure 2F),** and similarly found several compounds targeting nucleotide synthesis that displayed differential efficacy between RPMI and ftABS **(Figure 2F, G).** Together, these data suggest that altering environmental nutrient levels may particularly influence the response to compounds targeting cellular metabolism, but do not affect the efficacy of compounds that target cell intrinsic processes like signaling.

### Screening with a metabolism-oriented library identifies many compounds with differential efficacy in ftABS compared to RPMI

To further explore whether nutrient levels affect response to compounds targeting metabolic enzymes, we next asked whether a metabolism-focused library^30^ **(Figure 3A)** would identify more compounds exhibiting differential efficacy between RPMI and ftABS. Indeed, we found many additional compounds with differences in ability to impair proliferation of A549 cells in these two media **(Figure 3B, S3A).** Compounds targeting nucleotide metabolism, NAD^+^ metabolism, glutamine metabolism, and redox balance were particularly enriched **(Figure 3C-F),** with EC50 values of individual compound scoring that were significantly increased (e.g. GPP78, STF31) or decreased (e.g. azaserine, RSL3) by culture in ftABS, or were unable to be determined due to a complete lack of efficacy in ftABS (e.g. brequinar, FK866). These results further highlight the importance of nutrient levels when screening for anticancer compounds that target cell metabolism.

**Figure 3.**
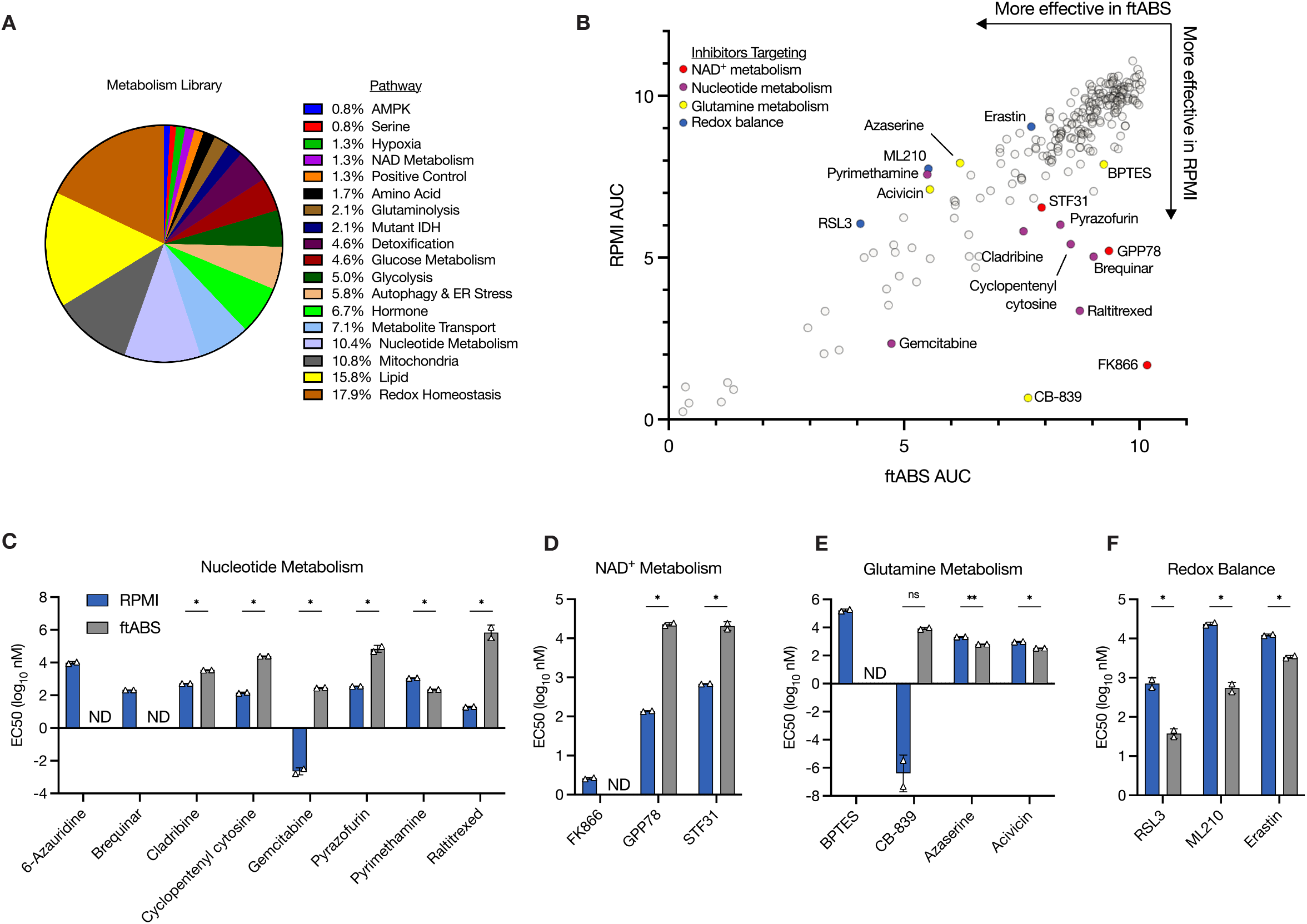
Hits for drugs with differential efficacy in standard versus serum-derived medium are enriched when a metabolism-oriented library is screened. **(A)** Breakdown of the Ludwig Metabolic Library 2 based on the molecular pathway associated with the annotated target of the compounds included in the library. **(B)** Plot of area under the curve (AUC) for A549 cells cultured in RPMI or ftABS and treated with the Ludwig Metabolic Library 2. Cells were treated for 72 hr with 10 doses of each drug up to a maximum dose of 20 μM, and dose response used to calculate AUC. Drugs in red target NAD+ metabolism proteins, drugs in purple target nucleotide metabolism proteins, drugs in yellow target glutamine metabolism, and drugs in blue target other redox metabolism proteins. **(C-F)** EC50 values for the indicated compounds targeting nucleotide metabolism (C), NAD^+^ metabolism (D), glutamine metabolism (E), or other redox metabolism proteins (F) as determined from the screen shown in 3B. Data shown are mean ± SD from two technical replicates (ND: an EC50 value could not be determined). Statistical test was performed using multiple unpaired t test with Benjamini and Hochberg FDR correction (ns, not significant; **q < 0.01; *q < 0.05).

### Physiologic nutrient levels cause resistance to NAMPT and DHODH inhibition by providing substrates for salvage pathways

Given that differences in nutrient levels between RPMI and ftABS likely drive the differential susceptibility to metabolism inhibitors, we sought to understand the mechanism by which this occurs for some of our top hits. FK866 is an inhibitor of NAMPT^31^, an enzyme that enables salvage of NAD^+^ from nicotinamide^32^, the NAD^+^ precursor that is abundant in RPMI **(Figure 4A).** We found FK866 to be highly effective in inhibiting proliferation of A549 cells cultured in RPMI, whereas cells cultured in ftABS were insensitive to FK866 treatment **(Figure 4B).** A549 cells were also protected from other inhibitors of NAMPT (e.g. GPP78, STF31) when cultured in ftABS **(Figure S3A).** We reasoned that salvageable NAD^+^ precursors other than nicotinamide may be present in biological fluids such as ftABS, but not RPMI, thus bypassing the need for NAMPT activity^32^. To test this hypothesis, we measured the concentration of nicotinic acid and found it to be 1 μM in ftABS **(Figure 4C).** Addition of nicotinic acid to this level in RPMI completely rescued the toxicity of FK866 in this medium **(Figure 4D)**. Another salvageable NAD^+^ precursor, nicotinamide mononucleotide (NMN), also rescued FK866 toxicity but was not detected in ftABS **(Figures S3B, 4C).**

**Figure 4.**
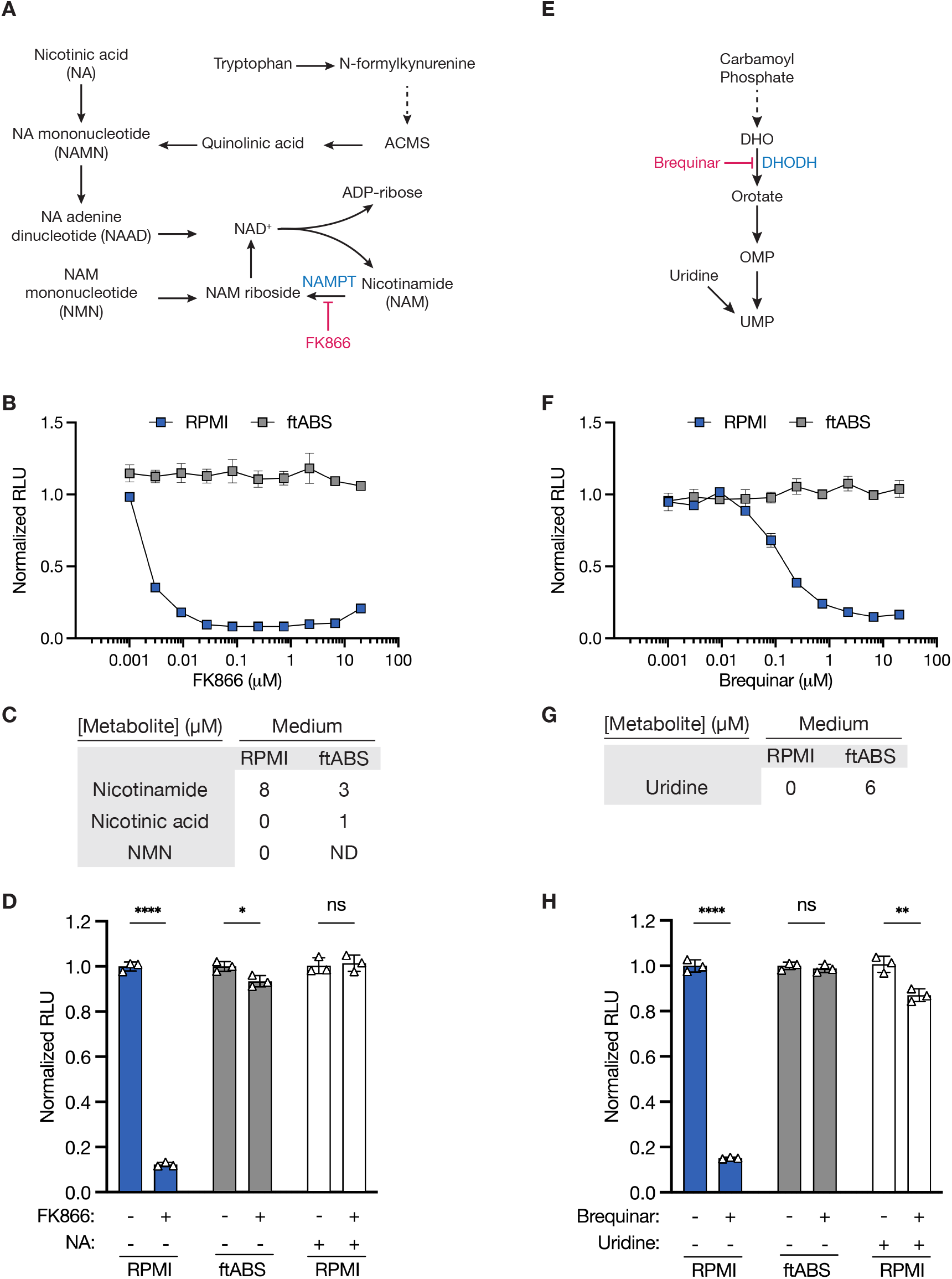
Physiologic metabolite levels can drive resistance to NAMPT and DHODH inhibitors. **(A)** Schematic illustrating the synthesis of nicotinamide adenine dinucleotide (NAD+) through the de novo Preiss-Handler or salvage pathways. NAMPT: nicotinamide phosphoribosyl transferase. FK866 is an inhibitor of NAMPT. **(B)** Effect of the NAMPT inhibitor FK866 on the number of A549 cells cultured in RPMI or ftABS for 72 hr as determined by CellTiter-Glo. Data shown are normalized to the vehicle-treated control and are the mean ± SD from two technical replicates. Data are from the screen in Figure 3B. **(C)** Concentrations of nicotinamide, nicotinic acid (NA) and nicotinamide mononucleotide (NMN) present in RPMI or ftABS. ND: measured but not detected. **(D)** Relative viability of A549 cells treated with vehicle or FK866 in ftABS or in RPMI, with or without addition of 1 μM NA for 72 hr as determined by CellTiter-Glo. Data are normalized to the vehicle-treated control and are the mean ± SD from three technical replicates. Statistical test was performed using multiple unpaired t test (ns, not significant; ****p < 0.0001; *p < 0.05). **(E)** Schematic illustrating the de novo synthesis and salvage pathways to generate the pyrimidine nucleotide uridine monophosphate (UMP). DHODH: dihydroorotate dehydrogenase. Brequinar is an inhibitor of DHODH. **(F)** Effect of the DHODH inhibitor brequinar on the number of A549 cells cultured in RPMI or ftABS for 72 hr as determined by CellTiter-Glo. Data shown are normalized to the vehicle-treated control and are the mean ± SD from two technical replicates. Data are from the screen in Figure 3B. **(G)** Concentration of uridine present in RPMI or ftABS. **(H)** Relative viability of A549 cells treated with vehicle or brequinar in ftABS or in RPMI, with or without addition of 6 μM uridine for 72 hr as determined by CellTiter-Glo. Data are normalized to vehicle-treated control and are the mean ± SD from three technical replicates. Statistical test was performed using multiple unpaired t test (ns, not significant; ****p < 0.0001; **p < 0.01).

Next, we found the compound brequinar, which inhibits DHODH in the *de novo* pyrimidine synthesis pathway **(Figure 4E),** to be more effective in A549 cells cultured in RPMI compared to ftABS **(Figure 4F).** Uridine is readily salvaged by most cells and can bypass the requirement to synthesize pyrimidines through DHODH^33^. We detected uridine in ftABS at a level that was sufficient to rescue the toxicity of brequinar observed in cells cultured in RPMI, as addition of uridine to these levels in RPMI led to brequinar resistance **(Figure 4G-H, S3C).** Together, these data highlight that the toxicity of some compounds in standard media can be explained by the absence of metabolites that are present in serum.

### Folate metabolism differs in cells cultured in serum-derived medium

Two inhibitors of one-carbon metabolism, methotrexate and pyrimethamine, were both more effective in slowing proliferation of cells cultured in ftABS **(Figure S3D).** This was surprising, given that cells are more resistant to methotrexate when cultured in media containing the physiological folate source 5-methyl tetrahydrofolate (5-MeTHF), rather than the non-physiological folate source present in RPMI, folic acid (FA)^34,35^. We quantified the levels of FA and 5-MeTHF in ftABS to be 2 nM and 10 nM, respectively, which are both lower than the concentration of FA present in RPMI **(Figure S3E).** We next cultured A549 cells in RPMI without folates for four days to deplete their intracellular folate reserves^34^, and then titrated back either 5-MeTHF or FA to determine their effect on cell proliferation in each medium. We found that folate-starved A549 cells failed to proliferate in either ftABS or RPMI lacking FA, but could proliferate in either medium if supplemented with FA or 5-MeTHF **(Figure S3F).** These data suggest that the folate levels present in ftABS are insufficient to support A549 cell proliferation following folate depletion, but confirm cells can proliferate in media with either FA or 5-MeTHF^34,35^. We next tested the efficacy of methotrexate on A549 cells that had been starved of folates and then cultured in media with either 5-MeTHF or FA, and indeed found that the cells were more sensitive to methotrexate when cultured in FA as previously reported^34,35^ **(Figure S3G).** Cells cultured in ftABS and FA were slightly more sensitive to methotrexate compared to cells cultured in RPMI and FA **(Figure S3G),** suggesting that additional nutrient differences between ftABS and RPMI may be contributing to the greater sensitivity to methotrexate in serum-based medium that warrants further study.

### Supraphysiological metabolite levels in standard medium can alter drug sensitivity

We found several inhibitors displaying differential efficacy between RPMI and ftABS that have been reported to impact the ferroptosis pathway of cell death^36^ **(Figure 5A-B, S4A).** Ferroptosis is characterized by an accumulation of toxic lipid peroxides that can arise from a block in cysteine import, glutathione depletion, or inhibition of the glutathione peroxidase GPX4^37^. RPMI contains supraphysiological levels of cystine **(Figure 5C),** and we found that supplementation of cystine to RPMI levels in ftABS rescues xCT and GPX4 inhibitor toxicity in ftABS **(Figure 5D, S4B).** Supplementation of cystine to RPMI levels in ftABS also re-sensitized A549 cells to CB-839 **(Figure 5D),** consistent with prior findings^4^.

**Figure 5.**
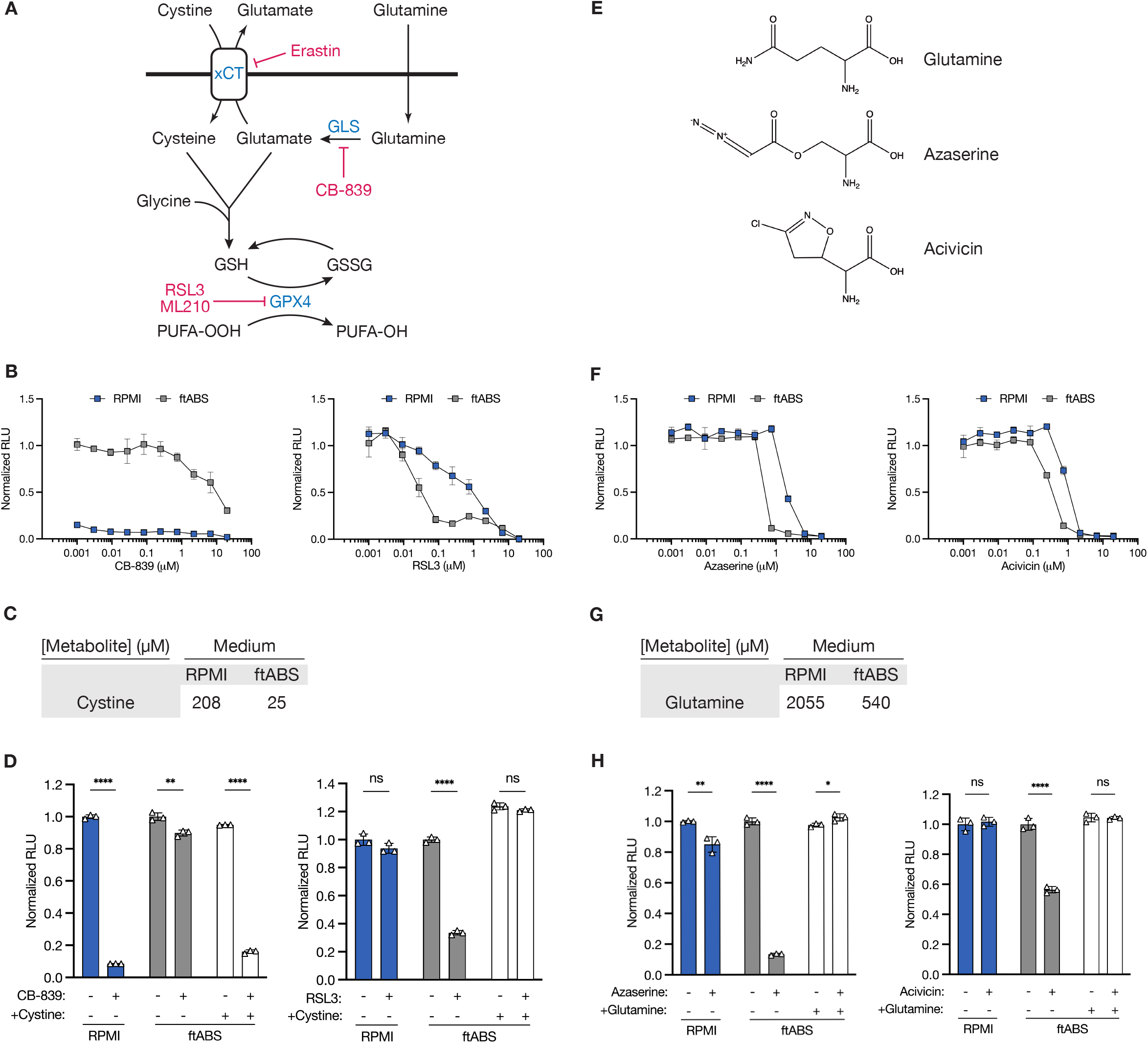
Supraphysiological metabolite levels in standard culture medium alters sensitivity to some compounds. **(A)** Schematic illustrating the synthesis pathways for glutathione (GSH) and for detoxification of lipid peroxides by glutathione peroxidase 4 (GPX4). xCT: cystine/glutamate antiporter, GLS: glutaminase, GSSG: glutathione disulfide, PUFA-OOH: peroxidated polyunsaturated fatty acid, PUFA-OH: polyunsaturated fatty acid. Erastin is an inhibit or of xCT. CB-839 is an inhibitor of GLS. RSL3/ML210 are inhibitors of GPX4. **(B)** Effect of CB-839 or RSL3 on the number of A549 cells cultured in RPMI or ftABS for 72 hras determined by CellTiter-Glo. Data shown are normalized to vehicle-treated control and are the mean ± SD from two technical replicates. Data are from the screen in Figure 3B. **(C)** Concentration of cystine in RPMI and ftABS. **(D)** Relative viability of A549 cells treated with vehicle, CB-839, or RSL3 in RPMI or in ftABS, with or without addition of 208 μM cystine (RPMI levels) for 72 hr as determined by CellTiter-Glo. Data are normalized to vehicle-treated control and are the mean ± SD from three technical replicates. Statistical tests were performed using multiple unpaired t test (ns, not significant; ****p <0.0001; **p < 0.01). **(E)** Chemical structures of glutamine and the glutamine analogs azaserine and acivicin. **(F)** Effect of azaserine or acivicin on the number of A549 cells cultured in RPMI or ftABS for 72 hr as determined by CellTiter-Glo. Data shown are normalized to vehicle-treated control and are the mean ± SD from two technical replicates. Data are from the screen in Figure 3B. **(G)** Concentration of glutamine in RPMI and ftABS. **(H)** Relative viability of A549 cells treated with vehicle, azaserine, or acivicin in RPMI or in ftABS, with or without addition of 2 mM glutamine (RPMI levels) for 72 hr as determined by CellTiter-Glo. Data are normalized to vehicle-treated control and are the mean ± SD from three technical replicates. Statistical tests were performed using multiple unpaired t test (ns, not significant; ****p <0.0001; **p < 0.01; *p < 0.05).

We identified two glutamine analogs, azaserine and acivicin **(Figure 5E),** that were more effective in slowing cell proliferation when added to cells cultured in ftABS compared to RPMI **(Figure 5F).** The toxicity of these compounds was reverted by addition of the supraphysiological level of glutamine present in RPMI to ftABS **(Figure 5G-H),** which has been reported to compete for uptake of these inhibitors into cells^38^. These data demonstrate that supraphysiological levels of metabolites in RPMI can protect cells from agents that may have greater activity when cells are exposed to more physiological nutrient levels.

Lastly, we tested whether the differential efficacy of select compounds targeting metabolic enzymes in affecting A549 cell proliferation was also retained across a panel of cancer cell lines. Indeed, consistent with data in A549 cells, we found that H1299, HCT-116, and HCC1806 cells were also more sensitive to FK866, brequinar and CB-839 when cultured in RPMI compared to ftABS **(Figure S5A-C),** and that azaserine was more effective when these cells are cultured in ftABS **(Figure S5D).** Interestingly, HeLa cells followed the same patterns, but did not exhibit differential sensitivity to CB-839 **(Figure S5A-D).** The differential response to erastin and RSL3 between media also varied across the cell lines tested **(Figure S5E-F),** further highlighting how screening in different nutrient conditions could be used to uncover biology that is not apparent from screening in standard conditions alone.

## DISCUSSION

To date, there are ~3,400 metabolites that have been reported as circulating in human blood, with many more detected but not quantified^24,39^. Yet, both standard culture media and synthetic blood-like media only capture a fraction of these metabolites and omit lipid species entirely in their base formulations. An advantage of ftABS is that this medium retains many small molecules present in serum at physiologic concentrations, including some lipids. However, the processing of ftABS to support high-throughput screening led to depletion of many lipids, since reducing protein content will remove lipoprotein and albumin, the main lipid carriers in the blood^40,41^. Removal of serum proteins may also influence drug response and cell metabolism, as exemplified by a recent study in which cancer cells experiencing ferroptosis via cystine starvation can be rescued by scavenging and breakdown of supplemented protein^42^, which can serve as a nutrient source^42,43^. Regardless, protein concentrations available to tumors may be lower than serum concentrations in some cases^44^, and conditions could be further optimized by adding back some components such as albumin or lipoproteins if relevant to a particular screen.

While screening in ftABS can uncover compounds whose mechanism of action depends on nutrient conditions, it may not always predict responses in animal models or patients as nutrient availability in tissues can differ from those found in the blood^23,45–47^. Regardless, the use of this screening approach to identify the relevant nutrient differences that impact response might still be informative for drug discovery. For example, many studies have proposed NAMPT as a cancer therapy based on the efficacy of NAMPT inhibitors in vitro, but efforts to target NAMPT for anticancer therapy ultimately failed in clinical trials due to lack of objective responses^48–51^, as well as dose-limiting toxicity in patients. While the activity of compounds in ftABS are unlikely to predict toxicity given inherent limitations of screening in vitro, screening in ftABS could highlight relevant modifiers of response to drugs. Indeed, determining which nutrients modify responses to a drug could be used to select contexts where responses are favored and contribute to patient selection. Additionally, identifying the relevant nutrient modifiers could suggest other dietary or pharmaceutical interventions to create metabolite conditions that may improve efficacy.

An inherent limitation of physiologic media including ftABS is that nutrient levels vary based on tissue location^23,45–47,31^, diet^52,53^, and genetics^54,55^, and no singular culture medium can recapitulate the diversity of conditions cells are exposed to in vivo. A particular limitation to ftABS as a culture media is that nutrient levels will vary across lots, although quantification of metabolite levels via mass spectrometry could still enable dissection of differential phenotypes that may be observed when using a new lot of ftABS for follow up work. However, preparing enough ftABS from the same lot to complete a screening study is an important practical consideration.

In the screens reported here, the majority of compounds with differential efficacy between media conditions showed higher activity in RPMI than in ftABS. This may be because the drug libraries screened were developed based on studies in standard culture conditions^27,56,57^. This inherent bias in the composition of some drug libraries may skew towards hits with activity in standard culture, and screening in ftABS or other physiological media conditions could be used as a tool to flag compounds with high activity driven by artificial media conditions. Considering that many novel small-molecule compounds are identified using phenotypic-based approaches^58,59^, comparing hits in standard culture with those found in ftABS may reduce pursuit of leads whose activity is driven by unique cell culture nutrient conditions.

Of the compounds displaying greater efficacy in ftABS, the glutamine analogs azaserine and acivicin are interesting given recent findings that treating mice with A549 xenografts with azaserine impaired tumor growth^60^. Clinical studies using glutamine analogs for advanced malignancies similarly showed promising responses dating back to 1950s, but dose-limiting systemic toxicities limited further development of these potential drugs^61^. To minimize systemic exposure, tumor targeted prodrugs of glutamine analogs were recently developed that are now in clinical trials based on robust responses in animal studies^61,62^ (ClinicalTrials.gov Identifier: NCT04471415).

Clinical success of anti-metabolite therapies clearly demonstrates that metabolism can be targeted for cancer treatment. However, progress to better match drugs targeting metabolism with the right patients has been hampered, in part, by limited utility of screening drugs in cancer cell lines under standard culture conditions to predict response in patients. There is also a perception that metabolism is too flexible for targeted therapy. Yet, recent evidence suggests that cancer cells experience constrained nutrient availability in tissues, and there may be an opportunity to selectively target cancer cell metabolism based on environmental nutrient availability. Therefore, screening approaches like the one highlighted here may help understand these physiological constraints and enable more effective targeting of cancer metabolism with both new and existing drugs.

## KEY RESOURCES TABLE

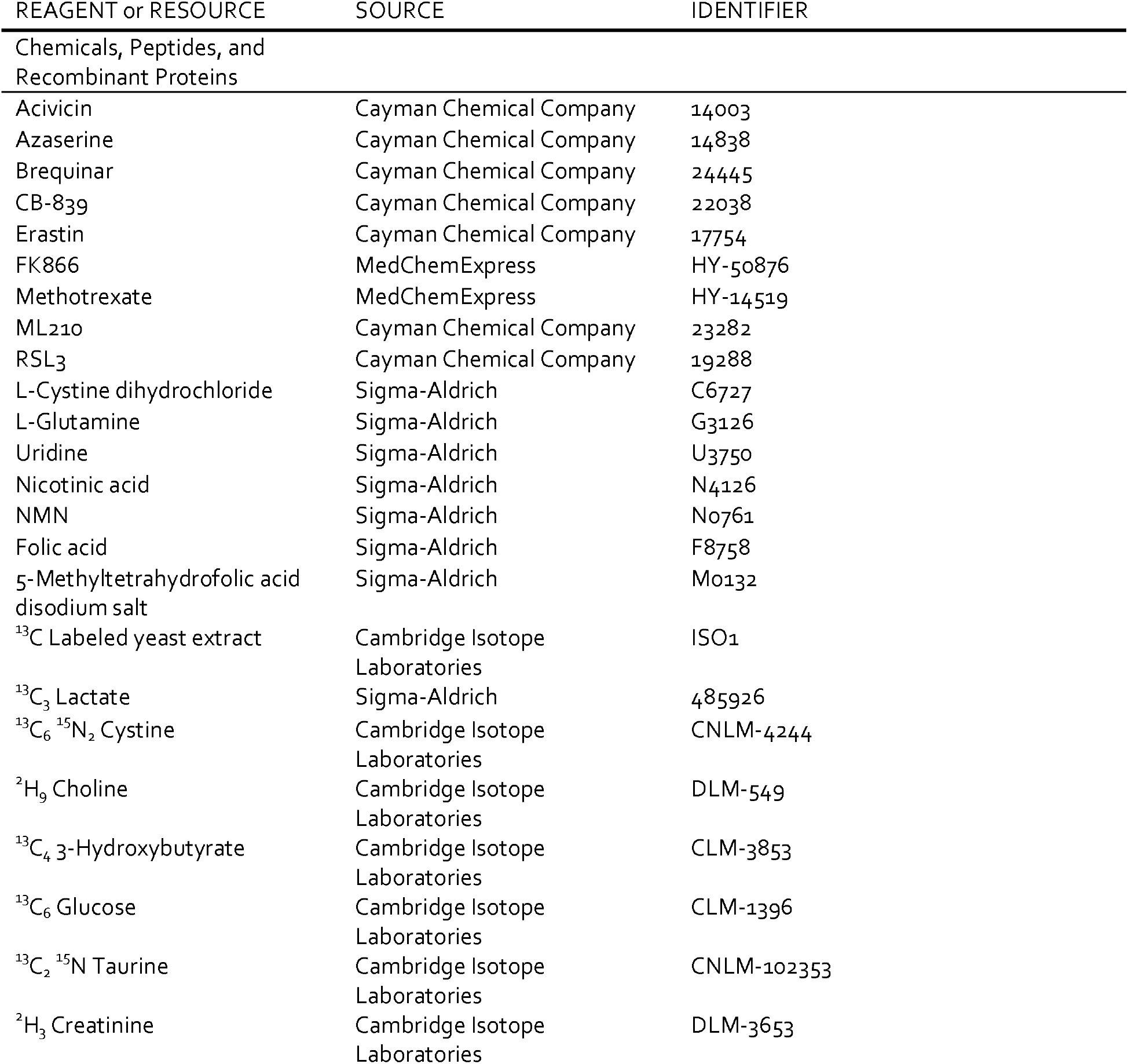
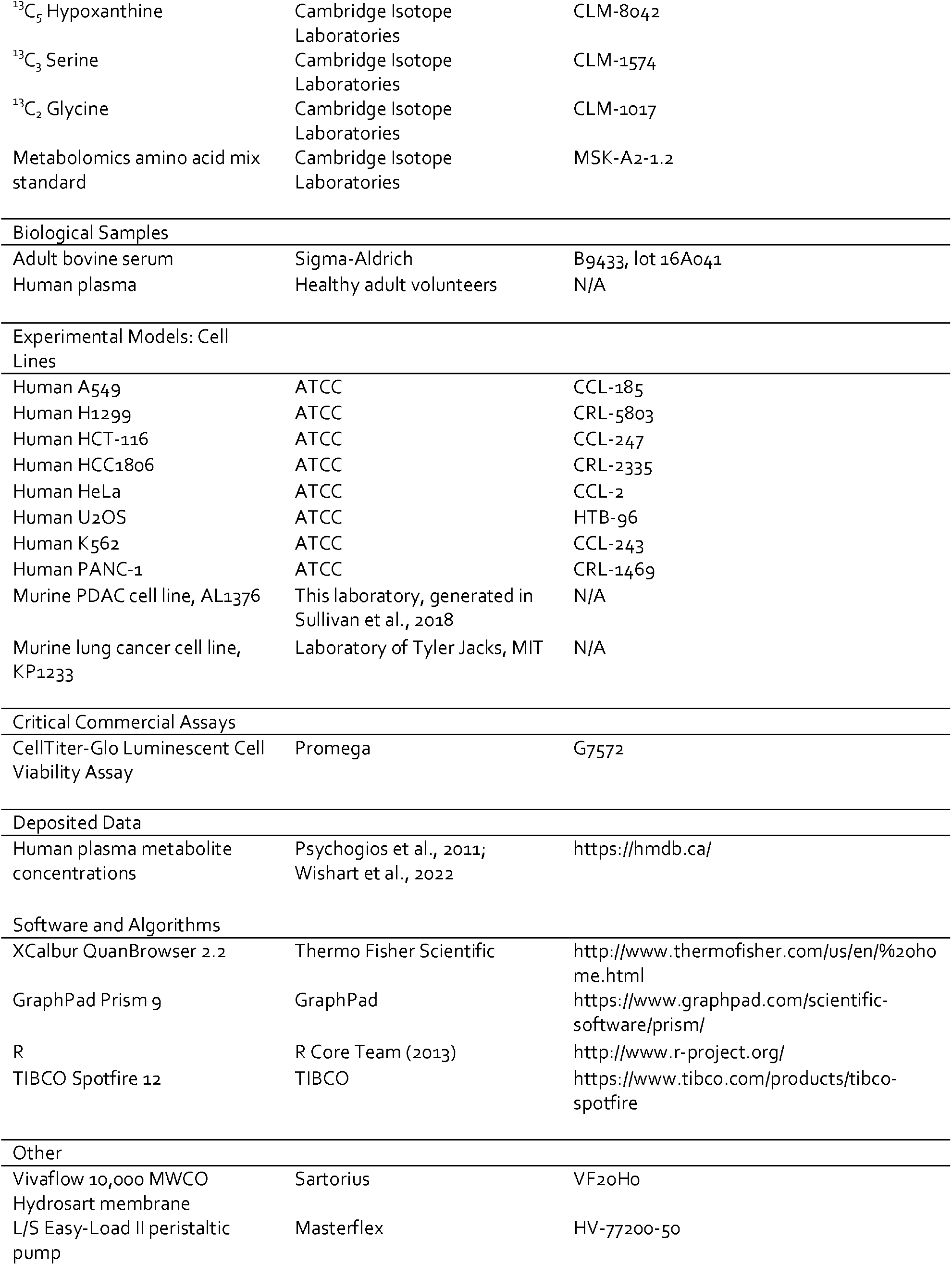

## CONTACT FOR REAGENT AND RESOURCE SHARING

Further information and requests for resources and reagents should be directed to and will be fulfilled by the Lead Contact, Matthew G. Vander Heiden.

## EXPERIMENTAL MODEL AND SUBJECT DETAILS

### Collection of human plasma

Ethical approval for the collection and analysis of human plasma was granted by the University of Cambridge Human Biology Research Ethics Committee (HBREC.17.20). Non-fasting blood samples from ten healthy adult volunteers (four female and six male, ages ranging from 22-40 years) were collected using 21G needles into 4 ml EDTA Vacutainer tubes (BD, 367839), then centrifuged at 800 xg for 5 min at 4°C to remove cells. Supernatants were then further centrifuged at 3,000 xg for 20 min at 4°C to remove platelets. Samples were snap-frozen and stored at −80°C prior to analysis by LC/MS analysis as described below. The time between collection and processing of each sample was <10 min.

### Cell lines and culture

Human cell lines used in this study were obtained from ATCC (A549: CCL-185; NCIH1299: CRL-5803; HCT-116: CCL-247; HCC1806: CRL-2335; HeLa: CCL-2; U2OS: HTB-96; K562: CCL-243; PANC-1: CRL-1469). Murine AL1376 cells were generated by the Vander Heiden lab previously^63^, and murine KP1233 cells were provided by Tyler Jacks (MIT). Lines were regularly tested for mycoplasma contamination using the MycoAlert PLUS Mycoplasma Detection Kit (Lonza BioSciences, LT07-710). All cells were cultured in a Heracell (Thermofisher, Waltham, MA) humidified incubator at 37°C and 5% CO_2_. Cell lines were routinely maintained in RPMI-1640 (Corning Life Sciences 10-040-CV) supplemented with 10% heat inactivated fetal bovine serum (Gibco 10437-028, lot 2372683RP), and for cell culture experiments, 10% dialyzed FBS (Gibco 26400-044, lot 2372678P) was supplemented, except where specified to have been cultured in ftABS or a related medium.

## METHOD DETAILS

### IDEXX panel

Blood chemistry of adult bovine serum, ftABS^B^, and dialyzed fetal bovine serum were analyzed in triplicate using the IDEXX Expanded Tox panel (IDEXX BioAnalytics). All samples were within assay range and were not diluted. Chemistries that were analyzed included total protein, albumin, globulin, glucose, cholesterol, HDL, LDL, triglycerides, sodium, chloride, potassium, calcium, phosphorus, TCO_2_ bicarbonate, BUN, creatinine, AST, ALT, ALP, CK, GGT, bile acids, and total bilirubin.

### Generation of flow-through adult bovine serum

100% adult bovine serum (Sigma-Aldrich B9433, lot 16A041) containing pen/strep was pumped through a Vivaflow 10,000 MWCO Hydrosart membrane (Sartorius VF20H0) according to manufacturer recommendations using a Masterflex L/S Easy-Load II peristaltic pump (Masterflex HV-77200-50) at 4°C. Filtrate containing metabolites (basal flow-through adult bovine serum, ftABS^B^) was collected until the sample reservoir reached approximately 50% of its starting volume. ftABS^B^ was then aliquoted and stored at −20°C. For cell culture experiments, except where indicated, 10% dialyzed FBS, 25 μM cystine, and 400 μM glutamine were also added to ftABS^B^ to generate complete medium (ftABS).

### Metabolomic analyses

#### Quantification of metabolite levels in biological fluids

Metabolite quantification in adult bovine serum, ftABS^B^, or human plasma samples was performed as described previously^23^. 5 μL of sample or external chemical standard pool (ranging from ~5 mM to ~1 μM) was mixed with 45 μL of acetonitrile:methanol:formic acid (75:25:0.1) extraction mix including isotopically labeled internal standards (see materials section). All solvents used in the extraction mix were HPLC grade. Samples were vortexed for 15 min at 4°C and insoluble material was sedimented by centrifugation at 16,000 x g for 10 min at 4°C. 20 μL of the soluble polar metabolite extract was taken for LC/MS analysis.

After LC/MS analysis, metabolite identification was performed with XCalibur 2.2 software (Thermo Fisher Scientific, Waltham, MA) using a 5 ppm mass accuracy and a 0.5 min retention time window. For metabolite identification, external standard pools were used for assignment of metabolites to peaks at given m/z and retention time, and to determine the limit of detection for each metabolite, which ranged from 100 nM to 3 μM (see Table S1 for the m/z and retention time for each metabolite analyzed). After metabolite identification, quantification was performed by two separate methods for either quantification by stable isotope dilution or external standard. For quantification by stable isotope dilution, where internal standards were available, we first compared the peak areas of the stable isotope labeled internal standards with the external standard pools diluted at known concentrations. This allowed for quantification of the concentration of labeled internal standards in the extraction mix. Subsequently, we compared the peak area of a given unlabeled metabolite in each sample with the peak area of the now quantified internal standard to determine the concentration of that metabolite in the sample. 57 metabolites were quantitated using this internal standard method (see Table S1 for the metabolites quantitated with internal standards). For metabolites without internal standards, quantification by external calibration was performed as described below. First, the peak area of each externally calibrated analyte was normalized to the peak area of a labeled amino acid internal standard that eluted at roughly the same retention time to account for differences in recovery between samples (see Table S1 for the labeled amino acid paired to each metabolite analyzed without an internal standard). This normalization was performed in both biological samples and external standard pool dilutions. From the normalized peak areas of metabolites in the external standard pool dilutions, we generated a standard curve describing the relationship between metabolite concentration and normalized peak area. The standard curves were linear with fits typically at or above r^2^ = 0.95. Metabolites which did not meet these criteria were excluded from further analysis. These equations were then used to convert normalized peak areas of analytes in each sample into analyte concentration in the samples. 40 metabolites were quantitated using this method.

#### Measurement of metabolite consumption and secretion

40,000 A549 cells were plated in 6-well plates in RPMI, allowed to attach overnight, washed twice with PBS, then treated with 2.2 or 6mL of RPMI or ftABS for 4,12, 24, 48, or 72 hr. At each time point, media was centrifuged at 845 x g for 3 min to remove cells, then flash frozen. 5 μL of media was mixed with 45 μL of HPLC-grade methanol:water (80:20) including 500 nM ^13^C/^15^N labeled amino acids (Cambridge Isotope Laboratories, MSK-CAA-1). Samples were then vortexed for 15 min at 4°C, centrifuged at 16,000 x g for 10 min at 4°C, and 20 μL of the soluble polar metabolite extract was taken for LC/MS analysis. Metabolite measurements were normalized to a labeled amino acid internal standard that eluted at roughly the same retention time. Metabolite concentrations were determined by normalizing to ion counts measured in fresh RPMI or ftABS media with known concentration values.

#### LC/MS analysis

Metabolite profiling was conducted on a QExactive bench top orbitrap mass spectrometer equipped with an Ion Max source and a HESIII probe, which was coupled to a Dionex UltiMate 3000 HPLC system (Thermo Fisher Scientific, San Jose, CA). External mass calibration was performed using the standard calibration mixture every 7 days. An additional custom mass calibration was performed weekly alongside standard mass calibrations to calibrate the lower end of the spectrum (m/z 70-1050 positive mode and m/z 60-900 negative mode) using the standard calibration mixtures spiked with glycine (positive mode) and aspartate (negative mode). 2 μL of each sample was injected onto a SeQuant^®^ ZIC^®^-pHILIC 150 x 2.1 mm analytical column equipped with a 2.1 x 20 mm guard column (both 5 mm particle size; EMD Millipore). Buffer A was 20 mM ammonium carbonate, 0.1% ammonium hydroxide; Buffer B was acetonitrile. The column oven and autosampler tray were held at 25°C and 4°C, respectively. The chromatographic gradient was run at a flow rate of 0.150 mL min^-1^ as follows: 0-20 min: linear gradient from 80-20% B; 20-20.5 min: linear gradient form 20-80% B; 20.5-28 min: hold at 80% B. The mass spectrometer was operated in full-scan, polarity-switching mode, with the spray voltage set to 3.0 kV, the heated capillary held at 275°C, and the HESI probe held at 350°C. The sheath gas flow was set to 40 units, the auxiliary gas flow was set to 15 units, and the sweep gas flow was set to 1 unit. MS data acquisition was performed in a range of m/z = 70–1000, with the resolution set at 70,000, the AGC target at 1×10^6^, and the maximum injection time at 20 msec.

#### LC/MS analysis of folate species

Extraction and LC/MS analysis of folate species was performed as previously described^34^. 10 μL of dialyzed FBS, ABS, or ftABS was mixed with 90 μL of extraction buffer(80:20 methanol: water with 2.5 mM sodium ascorbate, 25 mM ammonium acetate and 100 nM aminopterin). Samples were vortexed for 15 min at 4°C and insoluble material was sedimented by centrifugation at 16,000 x g for 10 min at 4°C. The supernatant was removed and dried under nitrogen. Samples were resuspended in 30 μL of HPLC-grade water, and 5 μL was injected onto an Ascentis Express C18 HPLC column (2.7 μm × 15 cm × 2.1 mm; Sigma-Aldrich). The column oven and autosampler tray were held at 30 °C and 4 °C, respectively. The following conditions were used to achieve chromatographic separation: buffer A was 0.1% formic acid and buffer B was acetonitrile with 0.1% formic acid. The chromatographic gradient was run at a flow rate of 0.25 ml min^-1^ as follows: 0–5 min where the gradient was held at 5% buffer B; 5–10 min at a linear gradient of 5–36% buffer B; 10.1–14.0 min at a linear gradient of 36–95% buffer B; and 14.1–18.0 min when the gradient was returned to 5% buffer B. The mass spectrometer was operated in full-scan, positive ionization mode. MS data acquisition was performed using three narrow-range scans: 438-450 m/z, 452-462 m/z and 470-478 m/z, with the resolution set at 70,000, the AGC target at 1×10^6^ and a maximum injection time of 150 ms. Quantification of folate species was performed with the Xcalibur QuanBrowser v.2.2 (Thermo Fisher Scientific) using a 5 ppm mass tolerance, and folate species were identified using chemical standards.

#### Untargeted metabolomics

5 μL of adult bovine serum or ftABS^B^ was mixed with 45 μL of HPLC-grade methanol:water (80:20) including 500 nM ^13^C/^15^N labeled amino acids (Cambridge Isotope Laboratories, MSK-CAA-1). Samples were vortexed for 15 min at 4°C and insoluble material was sedimented by centrifugation at 16,000 x g for 10 min at 4°C. 20 μL of the soluble polar metabolite extract was taken for LC/MS analysis. Untargeted metabolomics data were acquired in LC-MS mode with additional data-dependent MS/MS collected on pooled samples for identification of unknown metabolites. Data analysis was performed using Compound Discoverer 3.2 (Thermo Fisher Scientific) with an inhouse mass-list as well as online databases. Background compounds (i.e., compounds present in samples containing only extraction buffer or water) were excluded, as well as compounds with insufficient identification confidence. The median chromatographic peak area per group were used to generate a correlation plot.

#### Liquid chromatography-mass spectrometry lipidomics

Positive ion mode analyses of polar and nonpolar lipids were conducted using a liquid chromatography–mass spectrometry (LC-MS) system composed of a Shimadzu Nexera X2 U-HPLC (Shimadzu) coupled to an Exactive Plus orbitrap mass spectrometer (ThermoFisher Scientific). 2 μL of 100% adult bovine serum or ftABS^B^ were injected directly onto a 100 × 2.1 mm, 1.7-μm ACQUITY BEH C8 column (Waters). The column was eluted isocratically with 80% mobile phase A (95:5:0.1 v/v/v 10 mM ammonium acetate/methanol/formic acid) for 1 min followed by a linear gradient to 80% mobile phase B (99.9:0.1 v/v methanol/ formic acid) over 2 min, a linear gradient to 100% mobile phase B over 7 min, then 3 min at 100% mobile phase B. Mass spectrometry analyses were performed using electrospray ionization in the positive ion mode using full scan analysis over 220 to 1,100 m/z at 70,000 resolution and 3 Hz data acquisition rate. Other mass spectrometry settings were as follows: sheath gas 50, in source collision-induced dissociation 5 eV, sweep gas 5, spray voltage 3 kV, capillary temperature 300 °C, S-lens RF 60, heater temperature 300 °C, microscans 1, automatic gain control target 1×10^6^, and maximum ion time 100 ms. Lipid identities were determined on the basis of comparison to reference standards and reference plasma extracts and were denoted by the total number of carbons in the lipid acyl chain(s) and total number of double bonds in the lipid acyl chain(s).

### Cell proliferation

Cells were trypsinized, counted, plated in 2 mL complete RPMI at 10,000 cells per well of a 12-well plate, and allowed to attach overnight. Cells were then washed two times with 1 mL PBS per well, then treated with 2.5 mL of the indicated medium. A replicate plate was counted to determine starting cell number at the time of treatment. 72 hr post-treatment, cells were counted to calculate proliferation rate. Cell counts were determined using a Multisizer 3 Coulter Counter (Beckman Coulter) with a diameter setting of 10-30 μm. Proliferation rate was determined using the following formula:

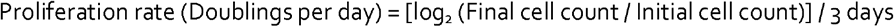

For assessment of proliferation by continuous live cell imaging, cells were trypsinized, pelleted and washed with PBS, counted, resuspended in the appropriate medium, then plated directly into clear 96-well plates at the cell densities as indicated. Plates were placed into an IncuCyte Live Cell Analysis Imaging System (Sartorius), and images were acquired every 3 hr using the 10x objective. Cell confluence was determined from a mask generated by the IncuCyte Zoom Analysis S3 2017 software’s standard settings.

### PRISM barcoded cell line screen

The PRISM multiplexed barcoded cell line assay was carried out as previously described^26^. 482 barcoded cancer cell lines were pooled together at equal representation and 33,000 cells plated into 6-well plates. After overnight attachment, cells were washed with PBS and medium was switched to 5 mL of RPMI or ftABS. One set of cells was then lysed and genomic DNA extracted to provide barcode representation at the start of the experiment. For remaining cells, medium was refreshed daily for a total of five days of growth, after which genomic DNA was extracted. Barcodes were then amplified from lysates^26^, and cell representation at the end of the experiment was calculated relative to starting representation (see Table S2 for log_2_FC values).

### High-throughput compound screening

#### Mechanism of Action Box (Moa Box) library

1536-well plates (Corning 3727BC) were pretreated with Novartis Institute of BioMedical Research MoA Box compound library^27^ using an Echo Acoustic Liquid Handler, dispensing 15 nL of each compound at doses ranging from 10.7 μM to 3.4 nM via 8-pt dose titration. 10 μM bortezomib was included as a kill control. A549 cells were trypsinized, resuspended in RPMI or ftABS, then plated in duplicate in 7 μL medium at 125 cells per well into compound-containing wells using a Multidrop Combi Reagent Dispenser (Thermo Scientific 5840320) plate dispenser. Cells were treated for 72 hr, after which 3 μL of CellTiter-Glo reagent (Promega, G7572) was added to each well, incubated at room temperature for 30 min, and luminescence read using a Pherastar plate reader. CellTiter-Glo measures intracellular ATP levels as a surrogate for viability. Activity values for each compound were normalized to DMSO-treated or bortezomib-treated wells, which served as 0% and 100% activity, respectively. The assays showed robust performance with RPMI Z□ = 0.65±0.04 and ftABS Z□ = 0.55±0.07.

#### FDA screen

A549 cells were trypsinized, resuspended in RPMI or ftABS, then 500 cells were plated in duplicate in 50 μL of medium per well of 384-well plates (Corning 3570BC) using an EL406 Washer Dispenser (BioTek). Cells were allowed to attach overnight, after which 50 nL of each compound from the SCREEN-WELL^®^ FDA approved drug library V2 (Enzo Life Sciences, BML-2843) was dispensed into each well using a Freedom Evo 150 Liquid Handler (Tecan) mounted with a 384W pin tool (V&P Scientific) at doses ranging from 10 μM to 1 nM via 5-pt dose titration. Cells were treated for 72 hr, after which 10 μL of CellTiter-Glo (Promega, G7572) reagent was added to each well, incubated at room temperature for 10 min, and luminescence read using Tecan M1000 plate reader. Luminescence values were normalized to DMSO-treated wells of the appropriate medium, and the area under the curve was quantified to assess the activity of each compound.

#### Metabolic library screen

A549 cells were trypsinized, resuspended in RPMI or ftABS, then 500 cells were plated in duplicate in 50 μL of medium per well of 384-well plates (Corning 3570) using a Multidrop Combi Reagent Dispenser (Thermo Scientific 5840320) plate dispenser. Cells were allowed to attach overnight, after which 100 nL of each compound from the Ludwig Metabolic Library 2 (provided to the ICCB-Longwood Screening Facility by the Ludwig Center at Harvard Medical School) was pin transferred into each well using a custom built Seiko Compound Transfer Robot (ICCB-Longwood Screening Facility, Harvard Medical School) at doses ranging from 20 μM to 1 nM via 10-pt dose titration. Cells were treated for 72 hr, after which 10 μL of CellTiter-Glo reagent (Promega, G7572) was added to each well, incubated at room temperature for 10 min, and luminescence read using a Perkin Elmer EnVision plate reader. Luminescence values were normalized to DMSO-treated wells of the appropriate medium, and the area under the curve was quantified to assess the activity of each compound. The assays showed robust performance with Z□ = 0.68±0.05 for cells treated with the positive control compound CB-839 in RPMI.

### Assignment of compounds targeting metabolism or signaling

A consensus metabolism gene list was generated by converging three gene lists generated in house (Muir) or previously published^28,29^ (Table S3). A list of signaling pathway genes was generated by downloading all relevant growth factor signaling or related pathways from KEGG (Table S3). Each compound from the MoA Box screen was assigned to a gene based on existing annotations^27^, and then designated whether it was included in the metabolism or signaling gene lists. The fraction of the library that targeted metabolism or signaling was then analyzed using z-score cutoffs, and significance was assessed via Fisher’s exact test.

### Drug enrichment analysis from MoA Box screen

For target enrichment calculations compounds were grouped into a set of actives with the desired profile (qAbsAC50 < 8 μM) and a set of inactives which did not show an effect in dose response in the A459 RPMI condition. Compounds from both sets were annotated with their published MoA targets^27^. The number of compounds with and without each target gene in the actives and inactives was used as input for a hypergeometric test to calculate p-values using the fisher.test function in R (R Core Team 2013) with alternative=“greater” settings. Multiple hypothesis adjustments were calculated with the p.adjust function in R with method=“BH”. Enrichment was calculated as (number of actives with target / number of actives without target) / (number of all cpds with target / number of all cpds without target).

### CellTiter-Glo assay

CellTiter-Glo assay for validation and rescue experiments was performed the same as in the high-throughput screens, with the following modifications: cells were plated into 96-well plates in 100 μL medium per well then allowed to attach for 24 hr; 100 μL of 2x treatment medium was added per well, and cells were incubated for 72 hr post-treatment; 40 μL of CellTiter-Glo (Promega, G7572) reagent was added per well, plates were mixed for 2 min using the linear shaking function of a Tecan Infinite 200 PRO plate reader (Tecan), then incubated for 10 min at room temperature before measuring well luminescence using Tecan plate reader. Luminescence values were normalized to vehicle-treated wells of the appropriate medium.

## QUANTIFICATION AND STATISTICAL ANALYSIS

All p values were calculated using Fisher’s exact test, multiple unpaired t test, ordinary one-way ANOVA followed by Dunnett’s multiple comparisons test, or Brown-Forsythe ANOVA followed by Dunnett’s T3 multiple comparisons test. All q values were calculated using two-tailed multiple unpaired t test and Benjamini and Hochberg FDR correction, or Brown-Forsythe ANOVA followed by Dunnett’s T3 multiple comparisons test. Welch’s correction was used in cases of unequal variance. GraphPad Prism 9 was used to calculate p and q values. All further statistical information is described in the figure legends.

## DATA AND SOFTWARE AVAILABILITY

### Data resources

Datasets can be found in Tables S1-4.

## ACKNOWLEDGMENTS

We thank Christopher Nabel, Alicia Darnell, Christopher Chidley and all members of the Vander Heiden lab for many useful discussions and experimental advice. We thank the NIBR Facilitated Access to Screening Technologies Lab, the Koch Institute High Throughput Sciences Facility, and the Harvard Medical School ICCB-Longwood Screening Facility for sharing their compound libraries and screening facilities. K.L.A. was supported by the National Science Foundation (DGE-1122374) and National Institutes of Health (NIH) (F31CA271787, T32GM007287). A.A received support as a Howard Hughes Medical Institute (HHMI) Medical Research Fellow. B.T.D. was supported by the NIH (F30HL156404, T32GM007753). R.F. was supported by the Knut and Alice Wallenberg Foundation (KAW 2019.0581). D.R.S. was supported by DF/HCC SPORE in Prostate Cancer Training Award (P50 CA090381), Harvard University KL2/Catalyst Medical Research Investigator Training award (TR002542), and Joint Center for Radiation Therapy Foundation. D.G. was supported by the Wellcome Trust (110302/Z/15/Z) and the NIHR Cambridge BRC. P.P.H. was supported in part by NIH (2T32CA071345-21A1). N.J.M. was supported by the Medical Research Council (MRC) (MR/T032413/1), NHS Blood and Transplant (WPA15-02), the Wellcome Trust (204845/Z/16/Z), and the NIHR Cambridge BRC. M.G.V.H. acknowledges support from the MIT Center for Precision Cancer Medicine, the Ludwig Center at MIT, Stand Up to Cancer, the Lustgarten foundation, an HHMI Faculty Scholars grant, and the NIH (R35CA242379, P30CA1405141).

## AUTHOR CONTRIBUTIONS

Conceptualization: K.L.A., A.A., A.M., D.S.A., M.G.V.H.; Methodology: K.L.A., A.A., D.C., J.H.C., C.K.S., D.R.S., P.P.H., L.M.G., A.M., D.S.A., M.G.V.H.; Investigation: K.L.A., A.A., D.C., B.T.D., R.F., J.H.C., C.K.S., A.D., T.K., D.R.S., S.R., S.E.H., M.W., S.H.C., T.T., D.G., C.W.N., C.J.Z., A.F., I.M.T.G., C.A.L., C.B.C., M.G.R., J.A.R., L.M.G., D.S.A.; Resources: N.J.M., C.A.L., C.B.C., M.G.R., J.A.R., D.S.A., M.G.V.H.; Writing – Original Draft: K.L.A., A.A., M.G.V.H.; Writing – Review & Editing: All authors; Supervision: D.S.A., M.G.V.H.; Funding Acquisition: M.G.V.H.

