## Supplementary figures and images for "Screening in serum-derived medium reveals differential response to compounds targeting metabolism"

### Supplemental Figures

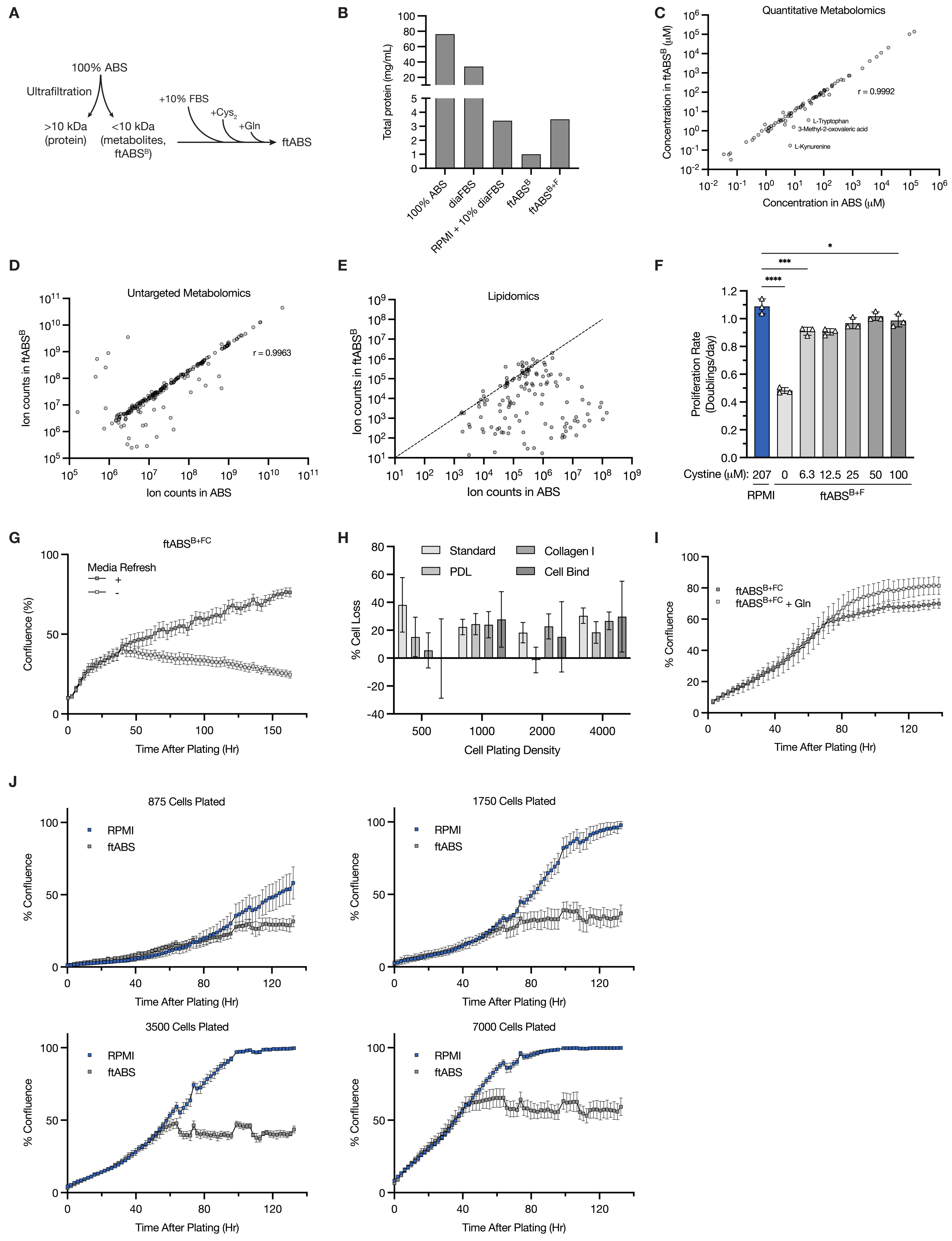

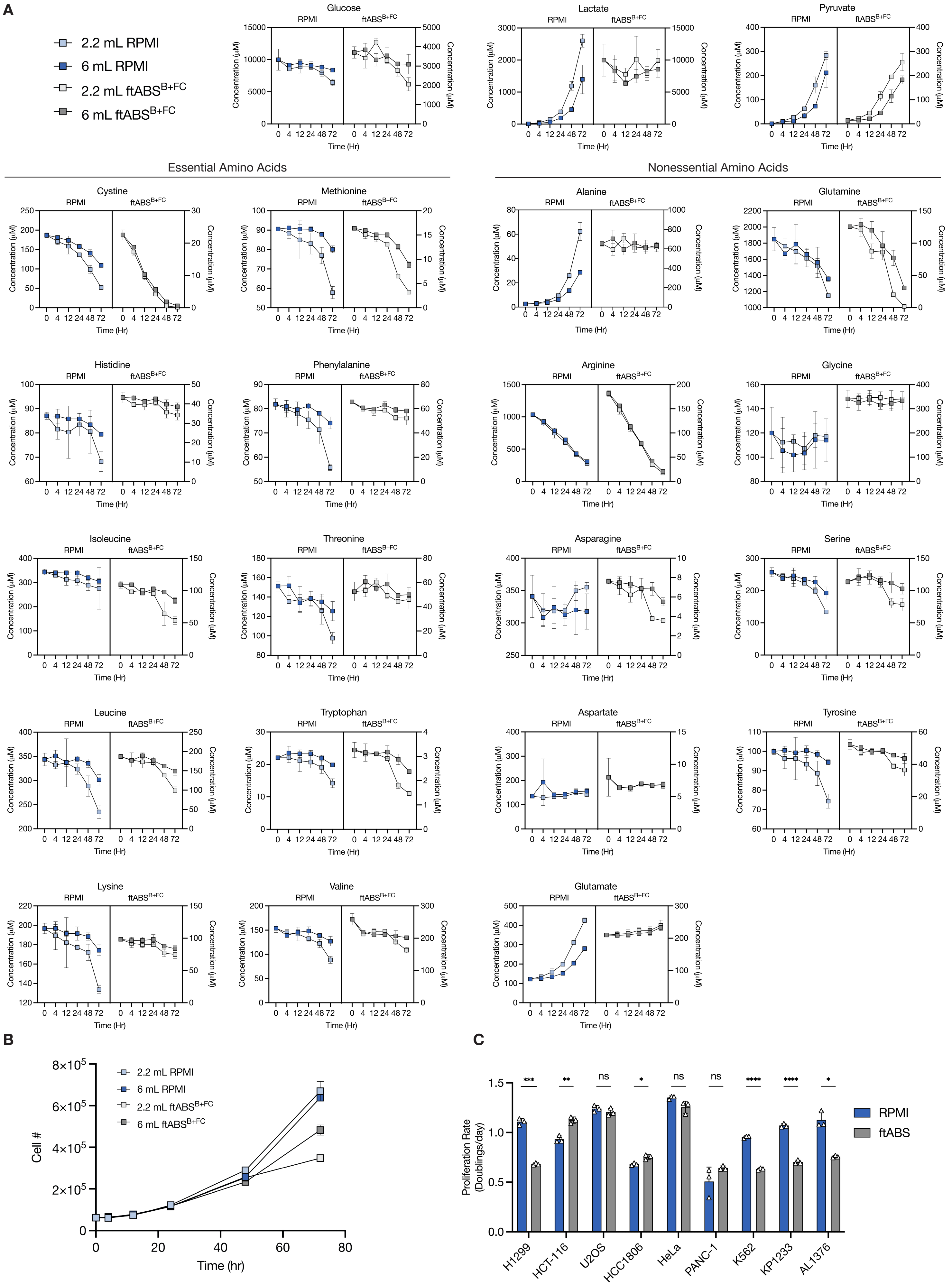

FIGURE S3

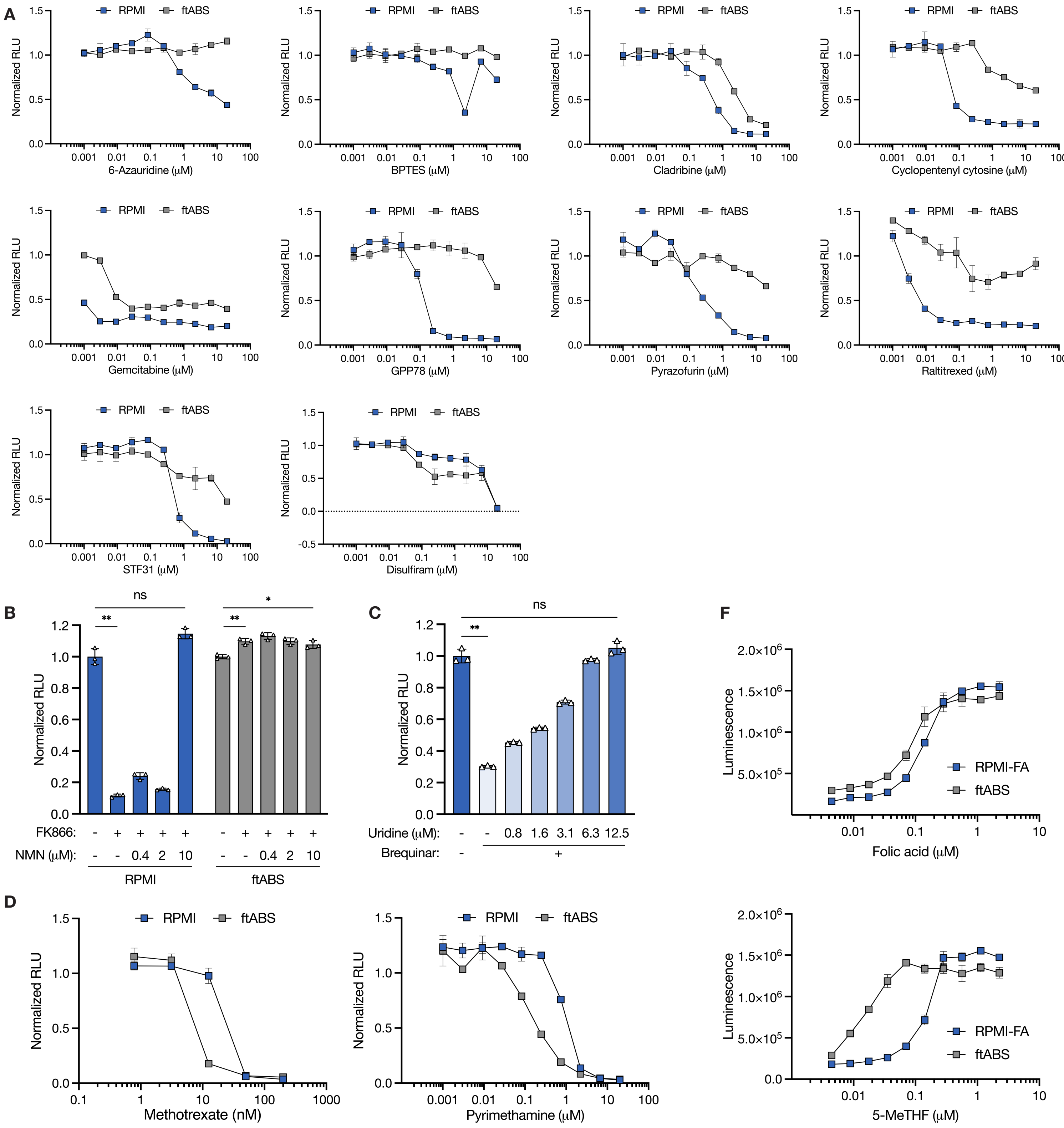


E

| [Metabolite] (µM) | Medium | | |
|---|---|---|---|
| | RPMI | ftABS | ABS |
| Folic acid | 2.27 | 0.002 | 0.002 |
| 5-MeTHF | 0 | 0.01 | 0.01 |

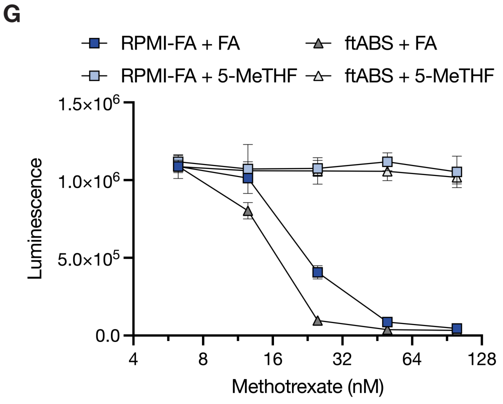

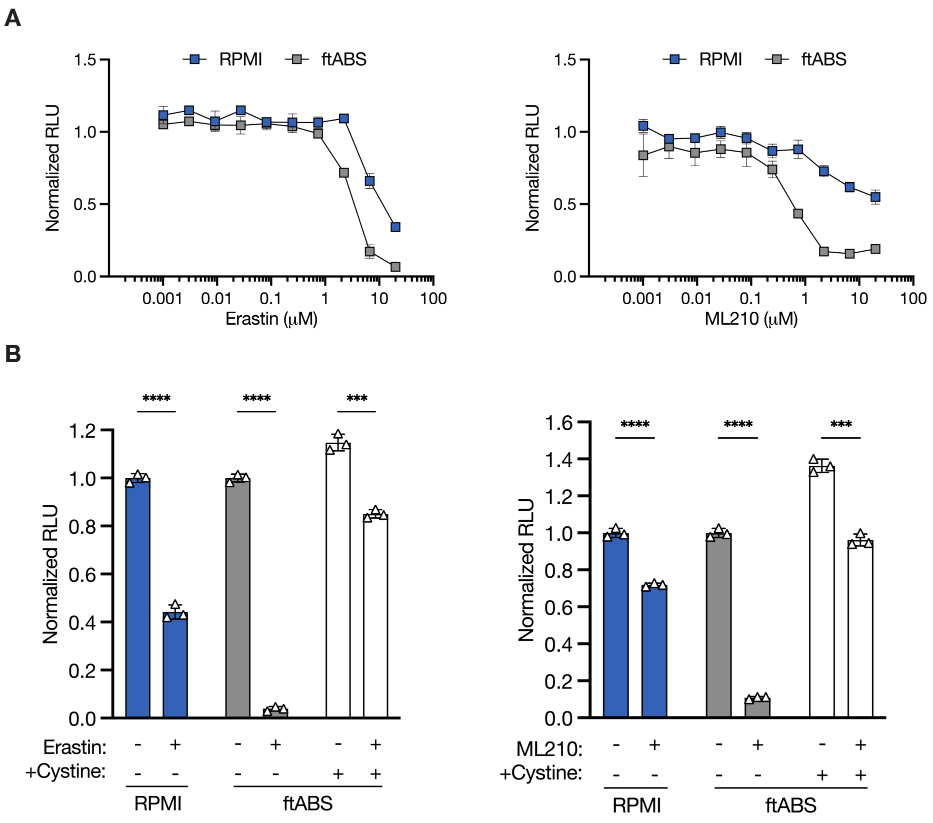

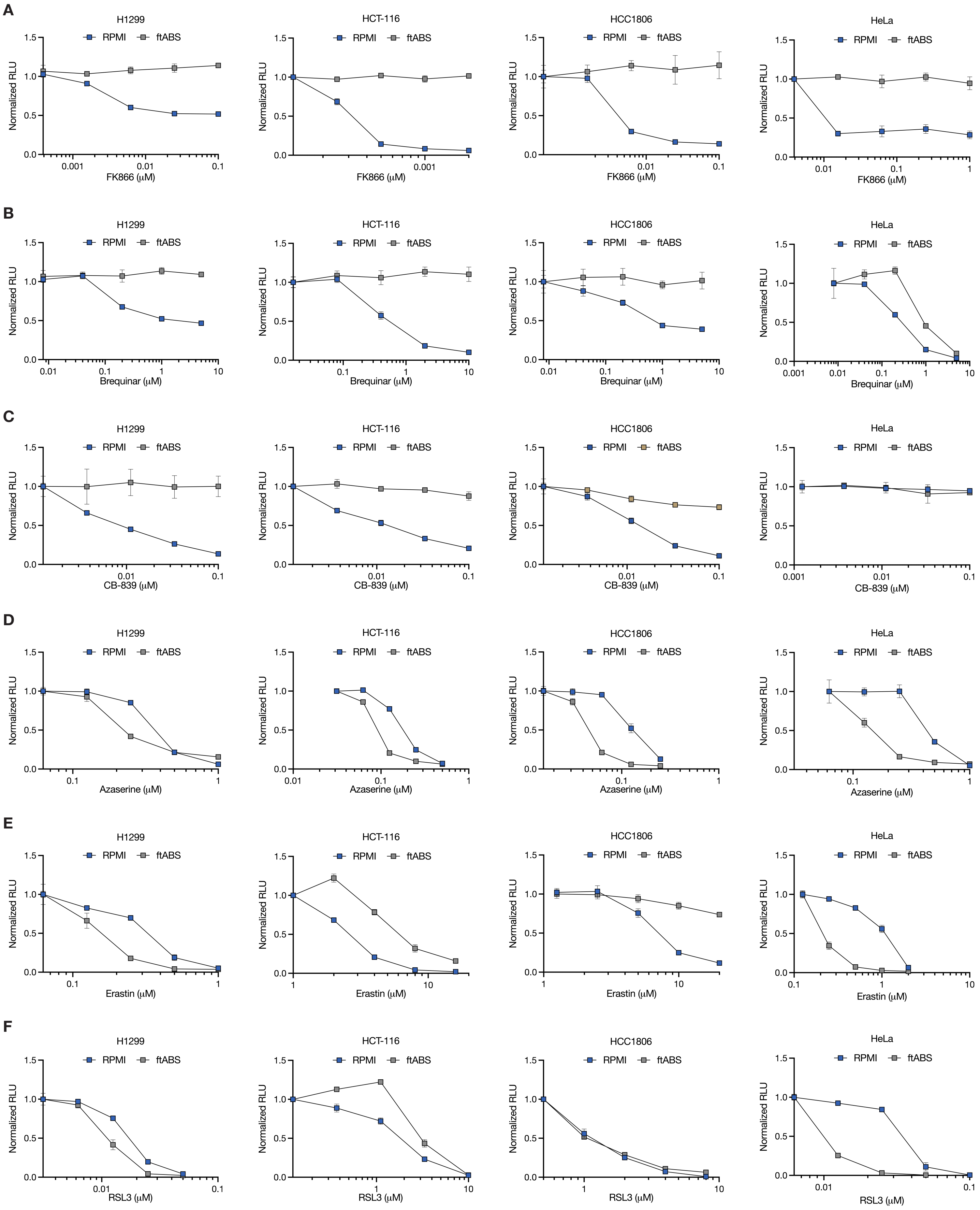
