## Supplementary Material for "Screening in serum-derived medium reveals differential response to compounds targeting metabolism"

**Supplementary Figure Legends**

**Figure S1. Optimization of a serum-based medium compatible with high-throughput screening.**

**(A)** Schematic depicting the generation of flow-through adult bovine serum (ftABS) from adult bovine serum (ABS). 100% ABS was processed via ultrafiltration through a 10 kDa membrane to generate basal ftABS (ftABS^B^), and the filtrate supplemented with 10% dialyzed fetal bovine serum (FBS), 25 μM cystine (Cys), and 400 μM glutamine (Gln) to generate the ftABS medium used in this study.

**(B)** Concentration of total protein present in 100% ABS, dialyzed FBS (diaFBS), RPMI with 10% diaFBS added, the ABS ultrafiltrate used to generate basal ftABS (ftABS^B^), and in basal ftABS after supplementation with 10% dialyzed FBS (ftABS^B+F^). Data shown are the mean ± SD from three technical replicates.

**(C-E)** Scatter plots of polar metabolite concentrations (n = 86 metabolites) (C), metabolite ion counts from untargeted metabolomics (n = 210 metabolites) (D), or lipid metabolite ion counts (n = 143 metabolites) (E) measured by LC-MS in ABS or basal ftABS (ftABS^B^). The values represent the mean of five (C & D) or four (E) LC/-MS measurements.

**(F)** Proliferation rate of A549 cells cultured for 72 hrs in RPMI or basal ftABS with 10% dialyzed FBS (ftABS^B+F^) supplemented with the indicated concentrations of cystine. Data shown are the mean ± SD from three technical replicates. Statistical test was performed using ordinary one-way ANOVA followed by Dunnett's multiple comparisons test (****p < 0.0001; ***p < 0.001; *p < 0.05).

**(G)** Percent confluence of A549 cells over time when cultured in basal ftABS with 10% dialyzed FBS and 25 μM cystine (ftABS^B+FC^), with or without refreshing the medium every 24 hr. Data are the mean ± SD from four technical replicates.

**(H)** Percent cell loss due to washing A549 cells twice with PBS. A549 cells were plated and cultured in RPMI at different densities in 384-well plates containing the indicated different attachment substrates. Cells were allowed to adhere overnight prior to washing and assessment of relative cell number as determined by CellTiter-Glo. Data are normalized to wells containing cells that were not washed and are the mean ± SD from 20 technical replicates. PDL: Poly-D-Lysine.

**(I)** Percent confluence of A549 cells over time when cultured in basal ftABS with 10% dialyzed FBS and 25 μM cystine (ftABS^B+FC^) with or without 400 μM glutamine supplementation. Data shown are the mean ± SD from ten technical replicates.

**(J)** Percent confluence of A549 cells over time when initially plated at densities of 875, 1750, 3500 and 7000 cells per well in a 1536-well plate, and cultured in RPMI or ftABS. Data shown are the mean ± SD from four technical replicates.

**Figure S2. Characterization of cell behavior to culture in ftABS.**

**(A)** Concentrations of glucose, lactate, pyruvate, essential amino acids, and nonessential amino acids measured in medium over time when A549 cells are cultured in 6-well plates in 2.2 or 6 mL of RPMI with 10% dialyzed FBS, or in basal ftABS with 10% dialyzed FBS and 25 μM cystine (ftABS^B+FC^), as indicated. Metabolite levels were measured by LC-MS and normalized to an internal standard, and metabolite concentrations were determined by normalizing to ion counts measured in fresh RPMI or ftABS media with known concentration values.

**(B)** Number of A549 cells over time when cultured in 2.2 or 6 mL of RPMI with 10% dialyzed FBS or basal ftABS with 10% dialyzed FBS and 25 μM cystine (ftABS^B+FC^). Data shown are from the same experiment in A, and are the mean ± SD from three technical replicates.

**(C)** Proliferation rates of the indicated human (H1299, HCT-116, U2OS, HCC1806, HeLa, PANC-1, K562) or mouse (KP1233, AL1376) cell lines when cultured in RPMI or ftABS for 72 hr. Data shown are the mean ± SD from three technical replicates. Statistical test was performed using multiple unpaired t test (ns, not significant; ****p < 0.0001; ***p < .001; **p < 0.01; *p < 0.05).

**Figure S3. Levels of specific nutrients dictate sensitivity to some drugs targeting metabolic proteins.**

**(A)** Dose-response curves of A549 cells treated with the indicated compounds from the screen shown in Figure 3B. Data shown are mean ± SD from two technical replicates.

**(B)** Relative viability of A549 cells treated with vehicle or FK866 in ftABS or in RPMI, with or without addition of nicotinamide mononucleotide (NMN) for 72 hr as determined by CellTiter-Glo. Data are normalized to the vehicle-treated control and are the mean ± SD from three technical replicates. Statistical tests were performed using Brown-Forsythe ANOVA followed by Dunnett's T3 multiple comparisons test (ns, not significant; **p < 0.01; *p < 0.05).

**(C)** Relative viability of A549 cells treated with vehicle or brequinar in RPMI, with or without addition of uridine for 72 hr as determined by CellTiter-Glo. Data are normalized to the vehicle-treated control and are the mean ± SD from three technical replicates. Statistical test was performed using Brown-Forsythe ANOVA followed by Dunnett's T3 multiple comparisons test (ns, not significant; **p < 0.01).

**(D)** Dose-response curves of A549 cells treated with methotrexate or pyrimethamine in RPMI or ftABS for 72 hr as determined by CellTiter-Glo. Data are normalized to the vehicle-treated control and are the mean ± SD from two (pyrimethamine) or three (methotrexate) technical replicates. Pyrimethamine data are from the screen presented in Figure 3B.

**(E)** Concentrations of folic acid or 5-methyltetrahydrofolate (5-MeTHF) present in RPMI, ABS, ftABS.

**(F)** Dose-response curves of A549 cells treated with indicated folate sources in RPMI or ftABS for 72 hr as determined by CellTiter-Glo. Cells were deprived of folates for four days prior to treatment. Data are the mean ± SD from three technical replicates.

**(G)** Dose-response curves of A549 cells treated with methotrexate and the indicated folate source (folic acid [FA] or 5-methyltetrahydrofolic acid [5-MeTHF]) in RPMI or ftABS for 72 hr as determined by CellTiter-Glo. Cells were starved of folates for four days prior to treatment. Data are the mean ± SD from three technical replicates.

**Figure S4. Cystine levels modulate sensitivity to compounds that induce ferroptosis.**

**(A)** Dose-response curves of A549 cells treated with erastin or ML210 in RPMI or ftABS for 72 hr as determined by CellTiter-Glo. Data are normalized to the vehicle-treated control and are the mean ± SD from three technical replicates. Data are from the screen in Figure 3B.

**(B)** Relative viability of A549 cells treated with vehicle, erastin, or ML210 in RPMI or in ftABS, with or without addition of 183 μM cystine for 72 hr as determined by CellTiter-Glo. Data are normalized to the vehicle-treated control and are the mean ± SD from three technical replicates. Statistical test was performed using multiple unpaired t test (****p < 0.0001; ***p < 0.001).

**Figure S5. The influence of nutrients on drugs targeting metabolism varies based on the cells tested.**

**(A-F)** Dose-response curves indicated cell lines treated with FK866 (A), brequinar (B), CB-839 (C), azaserine (D), erastin (E) and RSL3 (F) in RPMI or ftABS for 72 hr as determined by CellTiter-Glo. Data are normalized to the vehicle-treated control and are the mean ± SD from three technical replicates.

**Supplementary Tables 1-4**

Supplementary Table 1: Metabolite measurements.

Supplementary Table 2: PRISM barcode counts.

Supplementary Table 3: Metabolism and signaling pathway gene lists.

Supplementary Table 4: Screen data
